# Bayesian computation through cortical latent dynamics

**DOI:** 10.1101/465419

**Authors:** Hansem Sohn, Devika Narain, Nicolas Meirhaeghe, Mehrdad Jazayeri

## Abstract

Statistical regularities in the environment create prior beliefs that we rely on to optimize our behavior when sensory information is uncertain. Bayesian theory formalizes how prior beliefs can be leveraged, and has had a major impact on models of perception ^1^, sensorimotor function ^2,3^, and cognition ^4^. However, it is not known how recurrent interactions among neurons mediate Bayesian integration. Using a time interval reproduction task in monkeys, we found that prior statistics warp the underlying structure of population activity in the frontal cortex allowing the mapping of sensory inputs to motor outputs to be biased in accordance with Bayesian inference. Analysis of neural network models performing the task revealed that this warping was mediated by a low-dimensional curved manifold, and allowed us to further probe the potential causal underpinnings of this computational strategy. These results uncover a simple and general principle whereby prior beliefs exert their influence on behavior by sculpting cortical latent dynamics.

## Introduction

Past experiences impress upon neural circuits information about statistical regularities of the environment, which help us in all manners of behavior, from reaching for one’s back pocket to tracking a friend’s voice in a crowd and making inferences about others’ mental states. There is, however, a fundamental gap in our understanding of how behavior exploits statistical regularities in relation to how the nervous system represents past experiences. The effect of statistical regularities on behavior is often described in terms of Bayesian theory, which offers a powerful and principled framework for understanding the combined effect of prior beliefs and sensory evidence in perception ^1^, cognition ^4^, and sensorimotor function ^2,3^.

On the other hand, the effects of experience on neural activity have been described in terms of cellular mechanisms that govern the response properties of neurons ^5^. For example, natural statistics are thought to shape tuning properties and/or spontaneous activity of neurons through adjustments of synaptic connections in early sensory areas ^6–10^. Single-unit responses in higher-level cortices also encode recent sensory events ^11^, motor responses ^12–14^, and reward probabilities ^15–18^. However, an understanding of how experience-dependent neural representations enable Bayesian computations is lacking.

Recent studies have focused on an analysis of the geometry and structure of *in-vivo* cortical activity in trained animals and *in-silico* activity in trained recurrent neural networks (RNNs) to gain a deeper understanding of how neural dynamics might give rise to behaviorally-relevant computations ^19–27^. Following this emerging multidisciplinary approach, we analyzed the geometry of neural activity in the frontal cortex of monkeys and *in-silico* activity in RNNs in a Bayesian timing task, and found strong evidence that prior statistics establish curved manifolds of neural activity that cause the underlying representations of time to be biased in accordance with Bayes-optimal behavior.

## Task and behavior

We trained rhesus macaques to perform a time-interval reproduction task in which we could readily manipulate the prior belief and sensory uncertainty independently (Figure 1a). We refer to this as the Ready-Set-Go (RSG) task. Every trial was initiated by two fixation cues, a circle that the animal had to fixate and a square that instructed the animal to hold a joystick in its central position. While fixating, two visual flashes – Ready followed by Set – provided the first two beats of an isochronous rhythm. The animal was required to estimate the sample interval, *t_s_*, between Ready and Set (i.e., estimation epoch), and use this information in the subsequent production epoch to generate the omitted third beat (Go) by either initiating a saccade or moving the joystick to the left or right, depending on the location of a target cue in the periphery. Monkeys received reward if the produced interval, *t_p_*, between Set and Go was sufficiently close to *t_s_* (Figure 1b).

**Figure 1.**
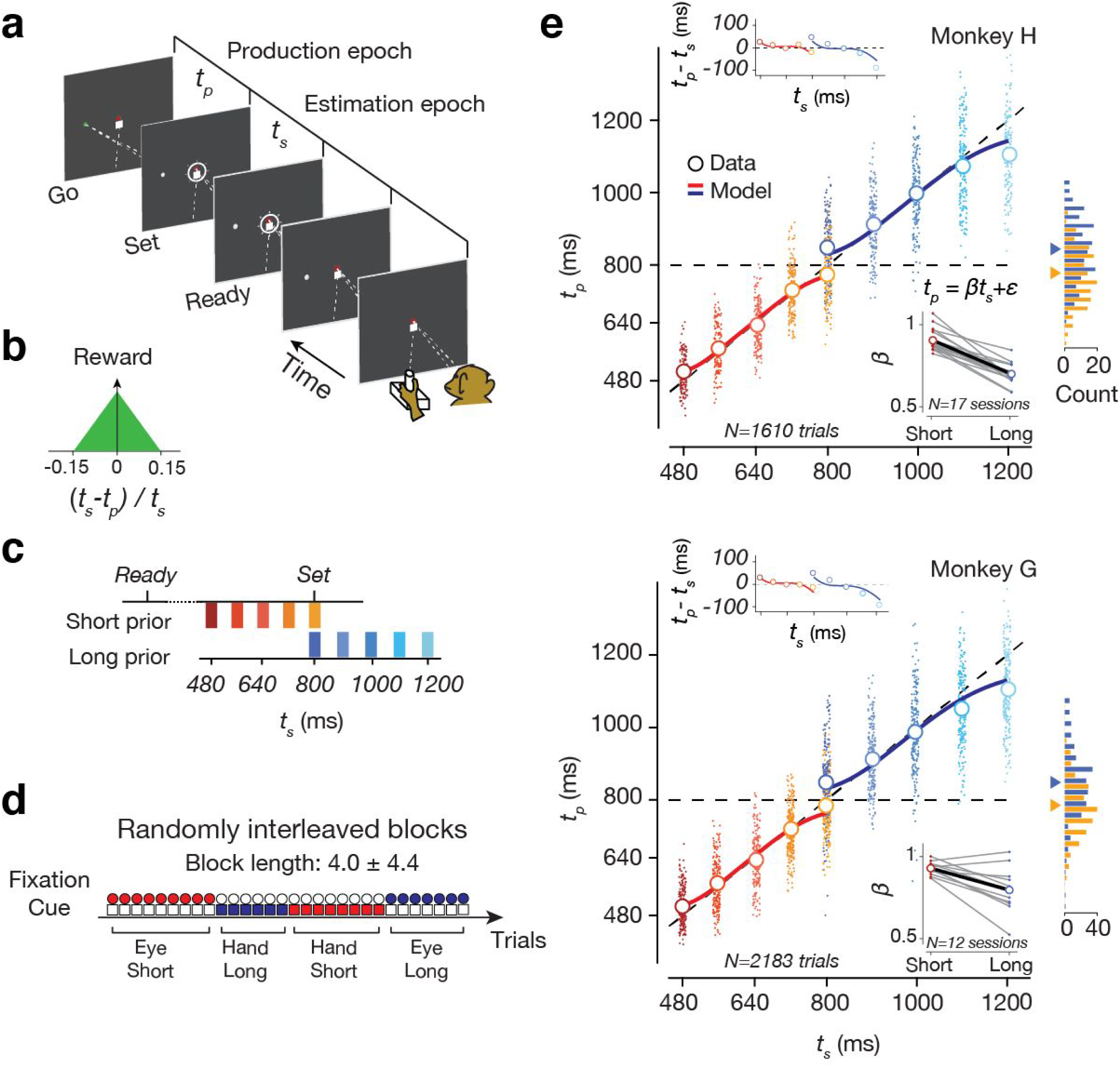
Task and behavior. a) Schematic of a single trial of the Ready-Set-Go task. A circle and a square fixation spot are presented at the center of the screen. The monkey fixates the circle and holds a joystick in center position. After a variable delay, a white target is presented to the left or right along the horizontal meridian. After another variable delay, a sequence of two flashes – Ready followed by Set – are presented around the fixation spot. The animal has to estimate the sample interval, *t_s_*, between Ready and Set (estimation epoch), and generate a delayed response toward the target either via a saccade or a movement of the joystick (production epoch). The produced interval, *t_p_*, between Set and movement initiation time (Go) has to match *t_s_*. b) Feedback. The monkey receives juice as reward (green region) if the relative error (*t_p_-t_s_*)*/t_s_* is smaller than *0.15*. Within this window, the amount of reward decreases linearly with the magnitude of error. At the time of feedback (i.e., immediately following the response), the target color changes to green or red in rewarded or non-rewarded trials, respectively. c) Prior distributions of *t_s_*. On each trial, *t_s_* is sampled from one of two discrete, uniform prior distributions (‘Short’ and ‘Long’) partially overlapping at *t_s_* = 800 ms. d) Trial types. The experiment consisted of 8 trial types: 2 prior conditions (Short and Long) x 2 effectors (Eye and Hand) x 2 target directions (Left and Right). The target direction was chosen randomly on a trial-by-trial basis. The 4 conditions associated with prior and effector were randomly interleaved across blocks of trials (see Methods for details). The block type was cued throughout the trial by the fixation spot: red circle and white square for Eye Short, red square and white circle for Hand Short, blue circle and white square for Eye Long, and blue square and white circle for Hand Long. e) Behavior. Top: A representative session showing individual *t_p_* values pooled across effectors and target directions (small filled circles) and corresponding averages (large open circles) for each *t_s_* for monkey H. The horizontal location of individual dots for each *t_s_* was jittered to facilitate visualization of individual *t* values associated with each *t_s_*. The red and blue lines are predictions based on fits of a single Bayesian model for both Short and Long prior conditions (see Methods and Figure S3). The diagonal shows the unity line. Right: Histograms of *t_p_* for the overlapping *t_s_* of 800 ms (horizontal dashed line) for each of the two prior conditions (Short: orange; Long: blue) with the corresponding averages (triangles). Top-left inset: Average error (i.e., bias) for each *t_s_* (data: circles; Bayesian model: solid lines). Bottom-right inset: Slopes of regression lines relating *t_p_* to *t_s_*. We used weighted linear regression to fit a line to individual data points for each prior condition separately (see Methods). Results for individual sessions is shown as small dots connected by gray lines, and the corresponding averages are shown as open circles connected by a black line. Bottom: The same as top for Monkey G.

To examine the neural basis of Bayesian integration, a critical aspect of the experimental design was that *t_s_* was sampled from one of two prior distributions, a ‘Short’ prior ranging between 480 and 800 ms, and a ‘Long’ prior ranging between 800 and 1200 ms (Figure 1c). Since the two prior conditions had an overlap at *t_s_* = 800 ms, the task offered a unique opportunity to characterize the representation of prior beliefs and how they might be integrated with ongoing sensory measurements. The full experiment consisted of eight conditions: two prior conditions (‘Short’ and ‘Long’), two response modalities (‘Eye’ and ‘Hand’), and two target directions (‘Left’ and ‘Right’). The prior condition and the desired effector switched across blocks of trials (block length: 4.0 ± 4.4 trials; uniform hazard) and were cued explicitly throughout every trial by the color of the fixation cues (Figure 1d). The target direction was chosen randomly across trials. The rationale for including two response modalities and two directions of response was to ensure that the neural correlates of Bayesian integration we identified would generalize across multiple experimental conditions.

To verify that animals learned to perform the task, we used a regression analysis to assess the dependence of *t_p_* on *t_s_* (Figure 1e, Figure S1). For both animals, the regression slope was positive in all conditions (Figure 1e, Table S1), demonstrating that their behavior correctly followed task contingencies. An important feature in both animals’ behavior was that the regression slopes were less than unity (Figure 1e, Table S1) indicating that animals systematically biased their responses toward the mean of the cued prior, consistent with Bayesian integration ^28^. In particular, the bias at the overlapping *t_s_* of 800 ms was in the opposite direction depending on the prior condition (rank-sum test, p<10^−43^ in animal H, p<10^−75^ in G; also see complementary analysis in Table S2). Importantly, this influence of prior on the bias was present immediately after block transitions, indicating that the animals were able to rapidly switch between priors using the cues (Figure S2). Bayes-optimal behavior additionally predicts that biases should be stronger for the Long prior condition for which measured intervals are more variable due to the scalar property of noise in interval timing ^29^. Consistent with this prediction, we found that the regression slope for the Short prior was significantly larger than that for the Long prior (Figure S1, Table S1). Finally, in agreement with previous work on variants of the RSG task in humans and monkeys ^28,30,31^, we found that behavioral statistics across animals, prior conditions and effectors were accurately captured by a Bayesian observer model (Figure S3, Table S3). Based on these results, we reasoned that the RSG task with two overlapping priors and various levels of measurement uncertainty is a suitable platform for investigating the representation of prior beliefs and the computational principles of Bayesian integration at the neural level.

## Electrophysiology

While animals performed the task, we recorded single-unit and multi-unit activity in the dorsomedial frontal cortex (DMFC; *N=617* and *741* in H and G, respectively) including the supplementary eye field (SEF), the dorsal region of the supplementary motor area (SMA), and pre-SMA. Our choice of recording areas was motivated by previous work showing a central role for DMFC in motor timing, movement planning and learning in humans ^32–37^, monkeys ^38–53^, and rodents ^54–60^.

During the estimation epoch, firing rates of single neurons were heterogeneous and exhibited rich dynamics that varied across experimental conditions (Figure 2a, see Figure S4 for more examples). A substantial proportion of neurons had distinct response dynamics depending on the prior condition (Figure 2a(i,iii,iv,v)). This observation is remarkable given that the two priors were switched rapidly across short blocks of trials. Indeed, knowledge about the prior condition altered neural responses at the very first trial after block transitions (Figure S5). We used a generalized linear model to quantify the degree to which spike counts of individual neurons during the support of the prior were modulated by elapsed time and the prior condition (see Methods). Results indicated that the activity of approximately 30% of neurons were modulated by time (27% Monkey H, 31% Monkey G; Figure 2c). As suggested by previous modeling studies, populations of neurons with such rich time-dependent responses may serve as a substrate for tracking elapsed time^61,62^. Nearly 60% of neurons changed their firing rate depending on the prior condition (65% Monkey H, 62% Monkey G; Figure 2c). The strong and systematic modulation of neural responses by the prior condition suggests that the neurons in this area were modulated according to the animal’s belief about the prior distribution of *t_s_*.

**Figure 2.**
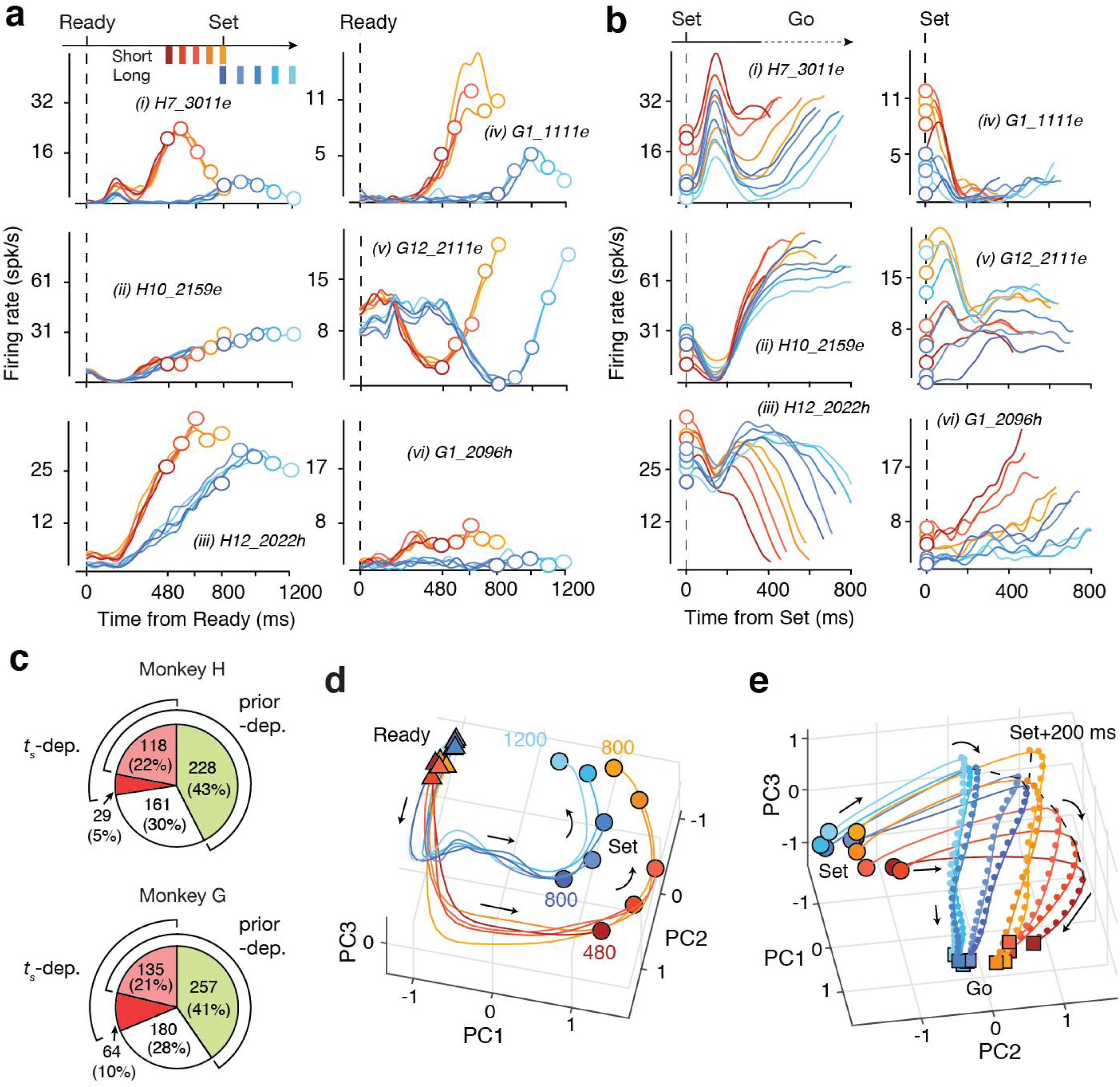
DMFC response profiles and neural trajectories. a) Firing rate of 6 example neurons during the estimation epoch labeled by Roman numeral (i-vi). Different shades of red and blue correspond to different *t_s_* intervals for the Short and Long prior conditions, respectively. Traces show activity from the time of Ready (vertical dashed line) to the time of Set (open circles), and the support of the prior is shown top left. Firing rates were obtained by smoothing averaged spike counts in 1-ms bins using a Gaussian kernel with a standard deviation of 25 ms. The label of each panel (e.g., H7_3011e) indicates the animal (H versus G) and the effector (e for Eye and h for Hand) associated with the traces. b) Firing rate of the same 6 neurons during the production epoch. Due to animals’ behavioral variability, production epochs for the same *t_s_* were of different durations. The plot shows the average activity of neurons from the time of Set (vertical dashed line) to the minimum *t_p_* for each *t_s_*. The color scheme is the same as panel a. c) A pie chart illustrating the proportion of neurons whose spike count during the prior support were dependent on the prior (“prior-dep.”) and/or *t_s_* (“*t*_s_-dep.”), determined by a generalized linear model (see Methods). The green region includes neurons that were only prior-dependent, the dark red, neurons that were only *t*_s_-dependent, light red, neurons that were both prior- and *t*_s_-dependent, white, the remaining neurons. d) Neural trajectories during the estimation epoch. A representative dataset is shown (Monkey H, Eye Left condition, see Figure S7 for other datasets). Trajectories are depicted in the subspace spanned by the first three principal components (PCs) in the estimation epoch using the same color scheme as in panel a. Triangles and circles represent the time of Ready and Set, respectively. Arrows illustrate the direction along which the trajectories evolve with time. e) Neural trajectories in the production epoch. Circles and squares represent the time of Set and Go, respectively (see Methods for how trials with different durations were handled). For each prior condition, the dashed line connects the neural states along the different trajectories 200 ms after Set. The small dots along each trajectory show neural states at 20-ms increments. The distance between consecutive dots is proportional to the speed at which activity evolves along a neural trajectory (e.g., higher speed for dark red compared to light blue).

Many DMFC neurons were also strongly modulated during the production epoch and exhibited temporally complex and heterogeneous patterns of activity (Figure 2b; see Figure S4 for more examples). Responses were often different at the time of Set because of prior- and *t*_s_-dependent modulations during the preceding estimation epoch. The presentation of Set was followed by transient modulations of firing rates, for about 200 ms (Figure 2b(i-iii,v)). Following this transient modulation, neurons exhibited a range of monotonic (e.g., ramping) or non-monotonic response profiles that were often organized systematically according to *t_s_* irrespective of the prior condition (Figure 2b(i-iii,v,vi)). A qualitative assessment indicated that responses of many neurons were temporally scaled with respect to *t_s_* (i.e., stretched in time for longer *t_s_*), an effect that was most conspicuous as a change of slope among the subset of ramping neurons (Figure 2b(ii,vi)). This temporal scaling is consistent with recent recordings in this area in a range of simple motor timing tasks^20,21,43,44,59^.

Several lines of evidence have led to the hypothesis that the relationship between neurons with such complex activity profiles and the computations they perform may be understood through population level analyses that depict the collective dynamics as neural trajectories governed by a dynamical system ^63–66^. Recent population-level analyses of neural activity in various higher cortical areas in a number of motor and cognitive tasks have provided support for this hypothesis ^19–23,67–71^. Following this line of work, we applied principal component analysis (PCA) to visualize the evolution of DMFC neural trajectories for various experimental conditions (see Methods). Our initial analysis indicated that neural responses associated with different effectors, target directions and epochs resided in different regions of the state space (Figure S6). Therefore, we applied PCA to trial-averaged neural responses across experimental conditions and task epochs separately.

For all datasets, the population activity in each epoch was relatively low dimensional: 3-4 principal components (PC) in the estimation epoch and 5-10 PCs in the production epoch explained nearly 75% of total variance (Figure S7). In the estimation epoch, neural trajectories associated with the two prior conditions were different at the time of Ready and became progressively more distinct throughout their evolution (Figure 2d; Movie S1). The most salient feature of population activity in this epoch was a rotation of neural trajectories that was temporally tuned to the support of the prior; i.e., approximately between 480 and 800 ms in the Short prior and between 800 and 1200 ms in the Long prior. The presence of rotational dynamics for the two priors was consistent with tuned responses of single neurons, many of which had nonlinear activity profiles that were specific to the support of the priors (Figure 2a(i,iii,iv)). Remarkably, these features were present in all experimental conditions (Figure S7) despite the fact that the corresponding neural activity patterns resided in different parts of the state space (Figure S6). These observations suggest that the rotational dynamics in DMFC may be the key for understanding the neural basis of Bayesian integration in the RSG task.

In the production epoch, consistent with observations of single neurons (Figure 2b), trajectories were at different initial states at the time of Set (Figure 2e; Movie S2). The Set flash caused a rapid displacement of neural states for nearly 200 ms. After the transient Set-triggered response, neural trajectories had an orderly structure with respect to *t_s_* and evolved toward a common terminal state (Go). A notable feature of neural trajectories in this epoch after the initial transient was an inverse relationship between *t_s_* and the speed with which responses evolved toward their terminal state. This effect was manifest in the displacement of neural states in 20-ms increments along neural trajectories associated with different *t_s_* intervals (Figure 2e). Specifically, neural trajectories appeared to evolve progressively slower for longer *t_s_*. Again, these features were expected based on the activity profile of single neurons (Figure 2b). The Set-triggered transient response was evident in the firing rate of single neurons (Figure 2b), and the change in speed was reflected in the temporal stretching of response profiles of many single neurons (Figure 2b (ii,iii,vi)). Both the role of speed in the control of movement initiation time ^21^, and the importance of initial state in adjusting the speed ^20^ have been demonstrated previously. The question that remains is how the brain utilizes a representation of the prior during the estimation epoch to set a suitable initial condition after Set so that the speed of the ensuing trajectories can take the information about the prior into account.

## Bayesian sensorimotor integration through latent dynamics

The common feature present across all experimental conditions in the state space was the rotation of neural trajectories during the support of the prior. How can a rotating trajectory encode prior belief and support Bayesian integration? An inherent property of a curved trajectory is that when it is projected onto a line connecting the two ends of the trajectory, equidistant points along the trajectory become warped. In other words, points near the ends of the projected line become biased toward the middle (Figure 3a), which is similar to the effect of Bayesian integration on the behavior (Figure 1e). This realization inspired the following hypothesis regarding how the rotation might serve as a substrate for Bayesian integration: neural states evolving along the rotating trajectory provide an implicit, moment-by-moment representation of the Bayesian estimate of elapsed time that could be decoded when projected onto a line in the state space.

**Figure 3.**
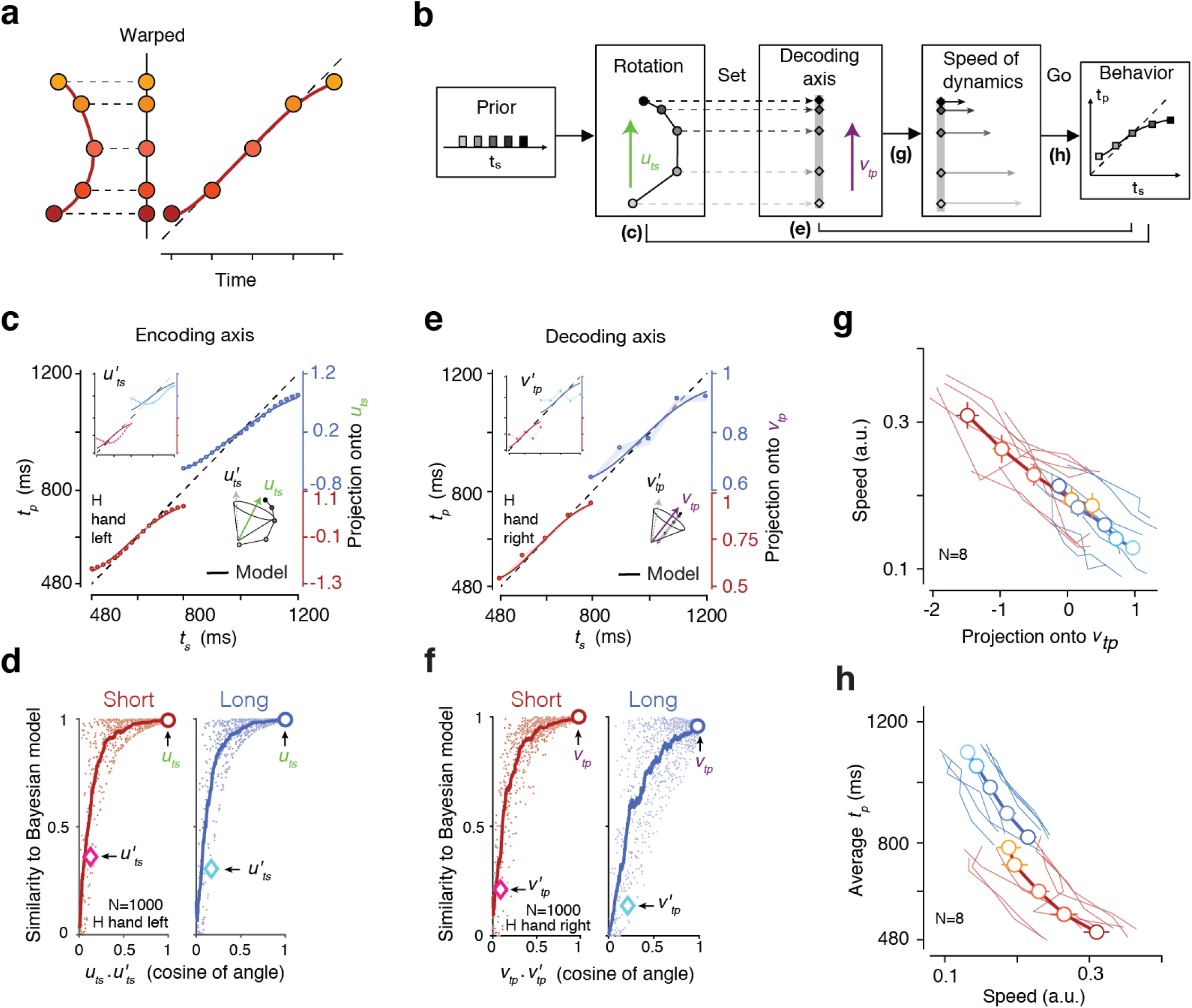
Neural signatures of Bayesian integration. a) A geometric illustration of how linear projection of points along a 2D curve onto a 1D line could cause warping mimicking the regression to the mean effect caused by Bayesian integration. b) The cascade of computations during estimation and production of the sample interval (*t_s_*) in the Ready-Set-Go task. The prior distribution of *t_s_* (leftmost panel) establishes rotational dynamics during the estimation epoch (second leftmost panel). Projection of the points along the rotating trajectory onto an encoding axis (green vector, *u_ts_*) creates a warped 1D representation of time that exhibits prior-dependent biases. Presentation of Set maps neural states onto a decoding axis (middle panel; purple vector, *v_tp_*). Neural states along the decoding axis serve as the system’s initial conditions during the production epoch. These initial conditions dictate the speed of neural trajectories (second rightmost panel) and allow the system to exhibit Bayes-optimal behavior (rightmost panel). The parenthetical labels (c), (e), (g) and (h) are evaluated quantitatively in the corresponding panels. c) Projection of neural states in the estimation epoch onto the encoding axis (*u*_*t*s_) as a function of *t_s_*. These projections yielded a warped representation of elapsed time whose relationship to actual elapsed time (abscissa) matched the prediction of a Bayesian model fit to behavior (line). The range of projections onto *u_ts_* (right ordinate axis) was linearly mapped onto the t_p_ range (left ordinate axis) for a meaningful comparison with the Bayesian fit. The plot shows a representative experimental condition (Monkey H, Hand Left condition). Circles show projections every 20 ms for Short (red) and Long (blue) prior conditions. Shaded areas represent 95% bootstrap confidence intervals (CIs). We tested other encoding axes (*u*’*_ts_*) within a cone centered on *u_ts_* (lower right inset) as shown for one random *u’_ts_* (top left inset) for which projected states (magenta for Short and cyan for Long) did not match the Bayesian predictions (line). d) A measure of similarity (based on R^2^: coefficient of determination) between neural states projected onto different vectors (*u*’_*t*_s) and the predictions of the Bayesian model as a function of the cosine of the angle between *u’_ts_* and the original *u_ts_* (Monkey H, Hand Left condition, see Figure S9 for other conditions). Small dots correspond to random *u’_ts_* vectors at various angles from *u_ts_* and lines are the respective moving averages. The circle and diamond symbols correspond to the original *u_ts_* and *u’_ts_* used for the top left inset of c). e) Projection of neural states 200 ms after Set onto the decoding axis (*V_tp_*). e) and f) show results of analyses on the decoding axis in the same format shown in c) and d) for the encoding axis. g) Speed at which neural states evolved during the production epoch (from Set + 200 ms to Go) as a function of the projection of the neural state at Set + 200 ms onto *v_tp_*. The speed was estimated by averaging distances between successive bins of the states in the state space. The thin lines correspond to individual datasets (2 animals x 2 effectors x 2 directions), and the thick line connecting circles show averages. Error bars are s.e.m. h) Average produced interval (*t_p_*) as a function of speed at which neural states evolved during the production epoch. Results are presented in the same format as in g.

To test this hypothesis, we asked whether projections of neural states during the support of the prior onto a one-dimensional ‘encoding axis’ could cause a regression to the mean consistent with Bayes-optimal behavior. Naturally, the answer depends on the choice of the encoding axis. Based on our understanding of the geometry of the problem (Figure 3a), we reasoned that a good candidate for the encoding axis was the vector pointing from the states associated with the shortest to the longest *t_s_* for each prior condition (*u_ts_;* Figure 3c). We projected neural states onto *u_ts_* to generate a one-dimensional representation of elapsed time during the support of the prior (Figure 3c). As predicted (Figure 3a), average neural states associated with each *t_s_* were warped with respect to the actual *t_s_* and exhibited biases toward the mean that matched the predictions of a Bayesian model fitted to the behavior (R^2^ = 0.993 for the Short prior, 0.996 for the Long prior; Figure 3c).

Our choice of *u_ts_* was motivated by an understanding of how projecting points along a curve onto a line causes warping (Figure 3a). To validate this choice, we tested other randomly chosen projection vectors (*u’_ts_*) within a cone in the state space that were up to 90 deg away from *u_ts_*. The similarity of projected states to the Bayesian model decreased progressively as a function of the angle between *u_ts_* and *u’_ts_* (Figure 3d), and for vectors far from *u_ts_*, projected states did not resemble the fits to the Bayesian model (Figure 3c, inset). These observations were consistent across priors, effectors and target directions (Figure S8). Together, these results suggest that the rotational dynamics in DMFC allow neural states to carry an implicit, continuous and instantaneous representation of the Bayesian estimate of elapsed time during the support of the prior.

We next asked how this implicit representation at the time of Set could influence *t_p_* at the end of the production epoch. Previous work has demonstrated that flexible production of timed intervals is made possible through adjustments of speed at which neural trajectories evolve toward an action-triggering state ^21,72–74^, and that this speed is determined by the initial conditions at the beginning of the production epoch ^20,72,75^. Accordingly, we evaluated the link between Set activity and *t_p_* in terms of a cascade of computations going from Set activity to initial conditions after Set, from initial conditions to speed of dynamics during Set-Go, and from speed to t_p_ (Figure 3b).

We hypothesized that the transient displacement of population activity following Set (Figure 2e) maps activity onto a “decoding axis” that serves as the initial condition, and that those initial conditions set the speed of the ensuing dynamics during the production epoch. An analysis of the structure of activity immediately after Set indicated that the Set-evoked transient response settled after nearly 200 ms (Figure S9). Therefore, we defined the decoding axis for each prior by a vector, *v_tp_*, that connected neural states associated with the shortest and longest *t_s_* 200 ms after Set (Figure 2e). As predicted by our hypothesis, average neural states projected onto *v_tp_* exhibited the characteristic regression to the mean present in the Bayesian model fits to the behavior (R^2^ = 0.993 for the Short prior and 0.951 for the long prior; Figure 3e). We also validated our choice of *v_tp_* over other decoding axes (*v’_tp_*) that were up to 90 deg away from *v_tp_* (Figure 3f) and found that our choice *v_tp_* yielded largest decoding efficiency and similarity with the Bayesian model.

Next, we asked whether the initial conditions along the decoding axis were predictive of the speed at which activity evolved afterwards. We estimated the speed as the average Euclidean distance (in the PC space accounting for at least 75% of the total variance) between neural states associated with successive bins (20 ms), divided by the duration of the production epoch. We then examined the relationship between speed and the projection of neural states onto *v_tp_*. Speeds were slower for the Long compared to Short prior condition, and for each prior condition, speed decreased monotonically with the initial conditions associated with longer *t_s_*, consistently across all conditions (Figure 3g; Pearson correlation, ρ_Short_=-0.77, p<10^−8^, ρ_Long_=-0.48, p<10^−2^). We verified that the relationship between speed and initial conditions held at the level of single trials (ρ_Short_=-0.12, p<10^−3^, ρ_Long_=-0.14, p<10^−4^). Corroborating previous findings ^20, 21, 72^, these results show that during flexible motor timing tasks, the brain utilizes initial conditions to adjust the speed of ensuing neural dynamics.

Finally, we verified that the speed of dynamics was predictive of the resulting *t_p_* across both priors and across all experimental conditions, both at the level of averages (Figure 3h; Pearson correlation, ρ_Short_=-0.62, p<10^−4^, ρ_Long_=-0.66, p<10^−5^) and single trials (ρ_Short_=-0.13, p<10^−3^, ρ_Long_=-0.14, p<10^−4^). Together, these step-by-step analyses support our prediction that the rotation during the estimation epoch supplies a Bayesian estimate of elapsed time that sets the speed of dynamics during the production epoch allowing animals to produce Bayes-optimal behavior.

## Trial-by-trial link between the encoding and decoding axes

To further substantiate the role of the encoding and decoding axes in Bayesian computations, we extended our analysis to single trials. An analysis of this kind is challenging since single-trial estimates of neural states can often be unreliable ^76^. Therefore, we focused on a behavioral session where we were able to record from a large number of neurons simultaneously (Monkey H: *N=48;* Figure S10 for monkey G) and thus could estimate momentary neural states with greater reliability. For this dataset, we first projected neural trajectories onto the subspace spanned by the first three PCs, which explained 80% of variance (Figure 4a top). We then computed projections of neural states onto *u_ts_* using a cross-validation procedure (see Methods), and generated distributions of projected values for each *t_s_* (Figure 4a bottom, Figure S10a top). To increase statistical power, we used standard scores (i.e., z-score) and combined data across effectors and directions. To evaluate the separation between distributions, we used a sensitivity index (d’) to measure the distance between each distribution to that associated with the mean *t_s_* (Figure 4b, Figure S10a bottom). This relative distance measure featured the two key properties of Bayesian integration. First, sensitivity curves for each prior exhibited a sigmoidal shape indicating that distributions associated with the shortest and longest *t_s_* were biased toward the mean *t_s_* for each prior (relative distances were significantly larger around middle *t_s_;* two-tailed paired t test, t(30)=3.56, p=0.001). Second, the overall distances were smaller for the long prior condition (two-tailed paired t-test on slope of regression, t(6)=3.91, p=0.008) consistent with the larger regression to the mean in this condition due to scalar variability. The difference between the two priors was also evident when we applied the same analysis to the decoding axis (two-tailed paired t-test on slope of regression, t(6)=4.08, p=0.007; Figure 4c,d, Figure S10b); i.e., relative distances were smaller for the long prior condition. These results provide compelling evidence that implicit representations along the encoding axis and explicit representations along the decoding axis were associated with the Bayesian estimate of *t_s_*.

**Figure 4.**
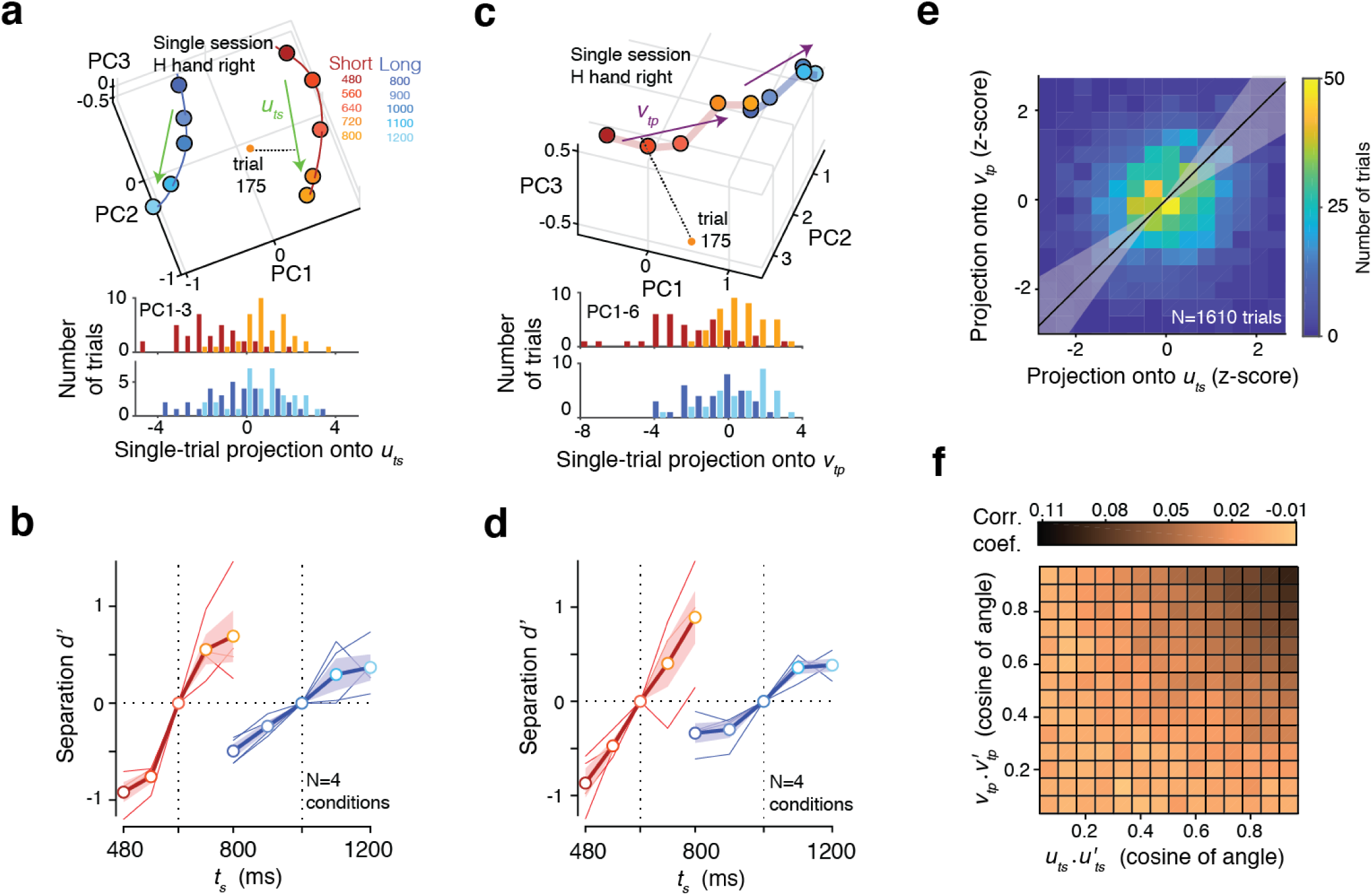
Bayesian computation at single-trial level. a) In a session with large number of simultaneously recorded neurons (Monkey H, Eye Right condition), we analyzed the distribution of projections onto *u_ts_* across single trials for each *t_s_*. Top panel shows the state space spanned by the first 3 principal components (PCs), the rotating neural trajectories within that space, and a single trial projected onto the corresponding *u_ts_* for each prior. Bottom panel shows the distribution of projected states for the shortest and longest *t_s_* of each prior condition in shades of red (Short) and blue (Long), respectively. Analyses were performed on cross-validated data (see Methods) within the subspace spanned by the first 3 PCs that explained 87.8% variance. (b) d’ measure quantifying the separation between the distribution of projected states for each *t_s_* to that for the middle *t_s_* (red for Short and blue for Long) as a function of *t_s_*. Thin and thick lines represent individual experimental conditions (2 effectors x 2 directions) and their corresponding average, respectively. c) In the same behavioral session, we analyzed the distribution of projection onto *v_tp_* across single trials for each *t_s_*. c) and d) show results of analyses on the decoding axis in the same format shown in a) and b) for the encoding axis. For the decoding axis, the distribution of projected states was computed in the subspace spanned by the first 6 PCs that explained 74.1% variance. e) Correlation between single-trial neural states at the time of Set projected onto *u_ts_* and the corresponding neural states 200 ms after Set projected onto *v_tp_*. To improve statistical power, we combined trials associated with different conditions (prior, effector, and direction) and different values of *t_s_* after z-scoring each dataset (line: best fit total-least-squares regression line; shading: 95% CI). f) Correlation coefficient between single-trial neural states projected onto *u’_ts_* and *v’_tp_* for 10000 randomly chosen pairs of *u’_ts_* and *v’_tp_*. The 2D histogram shows average correlations as a function of the cosine of angle between *u’_ts_* and *u_ts_* (abscissa) and between *v*’_*t*p_ and *v_tp_* (ordinate).

As a final assessment of representations in single trials, we asked whether neural states along the encoding axis (before Set) could be used to predict projections along the decoding axis (after Set). Remarkably, projections of neural states at the level of single trials onto *u_ts_* and *v_tp_* were significantly correlated (correlation coefficient: 0.118, p<0.001; 95% confidence interval from bootstrapping across trials: [0.070 0.170], Figure 4e, Figure S10c). This analysis demonstrates a systematic relationship between representations of time along the two axes. For example, when projections onto *u_ts_* for the shortest *t_s_* were closer to the expected state for a longer *t_s_* (due to trial-by-trial variability), the corresponding projections onto *v_tp_* were also biased in the same direction. Importantly, these trial-by-trial correlations were strongest for activity projected onto vectors *u_ts_* and *v_tp_* and decreased when activity was projected onto other random vectors (Figure 4f, Figure S10d). We also used the correlation strength to validate our choice of the temporal position of the decoding axis at 200 ms after Set. We found that correlations were strongest when *v_tp_* was computed from activity at around 200 ms after Set and decreased progressively when it was computed from earlier activity within the transient phase of the response (Figure S11). This result also rules out a trivial explanation of the correlation based on autocorrelation of firing rates near the time of Set. Together, these results suggest that the initial conditions along the decoding axis needed for Bayes-optimal behavior were computed by the rotational dynamics during the support of the prior.

## Recurrent network models of cortical Bayesian integration

Recurrent neural network models (RNNs) have proven useful in elucidating how neural populations in higher cortical areas support various motor and cognitive computations ^22,23,67,68,70,77,78^. They have also been useful for characterizing internally-generated dynamics in DMFC in flexible timing tasks ^20,21^. To gain further insight into how neural systems implement Bayesian inference, we trained RNNs to perform the two-prior RSG task (Figure 5a). On each trial, the network received the fixation cue as a tonic input whose value was adjusted by the prior condition. Afterwards, a second input administered Ready and Set via two pulses that were separated by *t_s_*. The network was trained to generate a linear ramping signal during Set-Go that would reach a fixed threshold (“Go”) at the correct time to reproduce *t_s_*. Using a suitable training strategy (see Methods), we were able to build RNNs whose behavior was accurately captured by a Bayesian observer model. In particular, responses were biased toward the mean for each prior condition, and biases were larger for the Long prior (Figure 5b).

**Figure 5:**
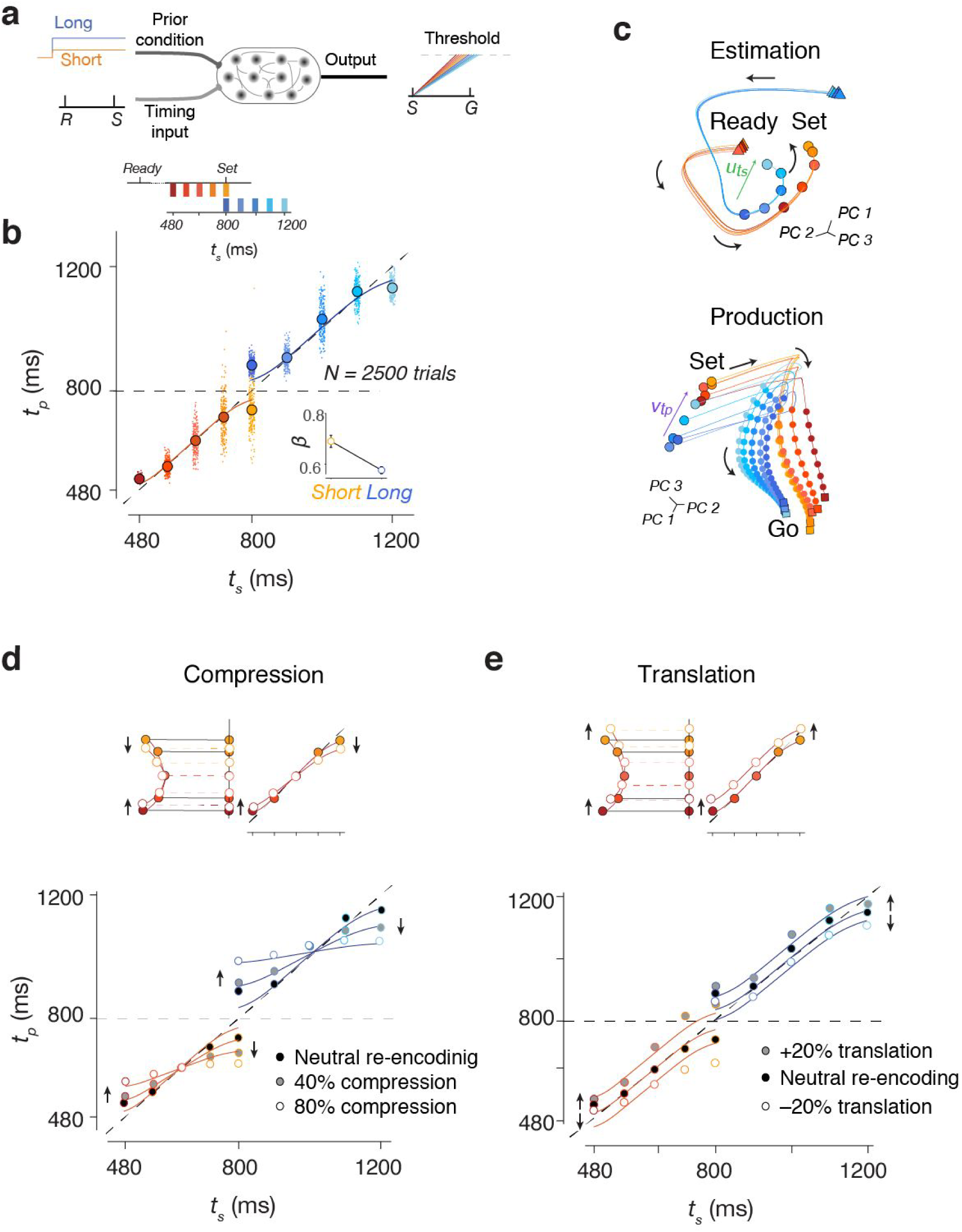
Recurrent neural networks re-creating Bayesian behavior and dynamics in cortical populations. a) Schematic of RNN experimental design. Prior condition is cued by tonic input (blue for long prior and orange for short). Second timing input provides the sample interval (*t_s_*) through Ready (R) and Set (S) pulses. The network was trained to generate a linearly ramping output between Set (S) and Go (G) whose slope was inversely related to *t_s_*. The produced interval (*t_p_*) was controlled by a threshold-crossing mechanism (dashed line). b) Network behavior shown using the same format as in Figure 1e except circles are filled to distinguish them from subsequent panels. Solid lines represent the behavior of the corresponding Bayesian ideal observer model. c) Network unit trajectories shown using the same format as Figure 2d,e. d) Top: Schematic showing the re-encoding process for compressive perturbation. Network states are compressed toward the state associated with the mean *t_s_*. Below: Network behavior with different levels of compressive re-encoding (gray: 40% compression; white: 80% compression) as well as neutral re-encoding with no effective perturbation (black; see Methods). Solid lines represent single parameter (w_m_) fits to the Bayesian model. e) Same as d for translational re-encoding toward larger (gray: 20% positive translation along the moving trajectory) and smaller (white: 20% negative translation against the moving trajectory) *t_s_*values. Solid lines represent the Bayesian model translated by an offset that was a single-parameter fit to the data using a least-squares procedure (w_m_ = 0.05).

Next, we examined activity in the trained RNNs. Like DMFC neurons, individual RNN units displayed heterogeneous response profiles and were strongly moduled during the support of the prior (Figure S12a,b). Similar to DMFC, the overall network activity was low dimensional during both the estimation and production epochs (Figure S12c,d). Most importantly, network population trajectories exhibited the same hallmarks of neural trajectories in DMFC. For instance, during the estimation epoch, unit trajectories exhibited rotational dynamics that were temporally tuned to the support of each prior (Figure 5c top), and during the production epoch, the initial condition and speed of trajectories were organized systematically depending on *t_s_* (Figure 5c bottom). To further assess the similarity between DMFC and RNN, we applied to the trained RNNs the same battery of analyses that was used to assess the cascade of computations in DMFC (Figure 3). Results indicated that the rotational dynamics established in the RNN resulted in a warped representation of *t_s_* along an encoding axis (*u_ts_*), which set the initial conditions along a decoding axis (*v_ts_*), and enabled speed-based dynamics leading to Bayes-optimal behavior (Figure S12 g,h). Based on these results, we concluded that the trained RNNs provided a suitable platform for validating and further delving into the importance of rotational dynamics in Bayesian computations.

A key advantage of training RNNs was that it allowed us to move beyond correlational observations relating rotational dynamics to Bayesian computation, and establish a causal link between the two. A major challenge in performing causal experiments on low-dimensional population activity is to successfully orient the perturbation along computationally-relevant dimensions of neural activity ^23,79,80^. Such targeted-dimensionality perturbation experiments are currently not feasible *in-vivo*, but they are *in-silico*. Accordingly, we developed an *in-silico* perturbation strategy, in which we halted the RNN shortly before Set, altered its state and then released the network to evaluate the outcome of the perturbation on the behavior. Importantly, the perturbation systematically targeted projections of neural states onto the encoding axis – a strategy that we refer to as re-encoding (Methods). We reasoned that if the rotational dynamics and the corresponding *u_ts_* are causally involved in creating biased neural representations, perturbing network state along this axis would lead to predictable behavioral outcomes.

Using this strategy, we perturbed network activity just preceding the time of Set in two ways: 1) compression around the middle *t_s_* (mean of the prior) along *u_ts_* and 2) linear translation along *u_ts_*. According to our hypothesis, the projection of activity along *u_ts_* provides an implicit representation for the Bayesian estimate of *t_s_*. This hypothesis makes specific predictions for how the compressive and translational perturbations would impact the behavior. The compression should lead to increased bias toward the mean *t_s_* (Figure 5d). The translation, on the other hand, should result in a translation in the value of *t_p_* towards longer or shorter intervals (Figure 5e) depending on the direction of the translation. Results confirmed these predictions: *t_p_* values exhibited progressively larger regression to the mean for larger compressive perturbations (Figure 5d), and underwent an overall upward or downward shift as a result of translation (Figure 5e). These causal experiments provide additional evidence that the brain might also be using its prior-dependent rotational latent dynamics to implicitly represent the Bayesian estimate of *t_s_*.

## Discussion

The classic formulation of Bayesian models assumes that the observer integrates a sensory likelihood function with the prior probability distribution to compute a posterior distribution, and applies a cost function to the posterior to extract an optimal estimate. Inspired by this formulation, theoretical and experimental studies have sought to find representations of various components of Bayesian inference at the level of single neurons. For example, some studies have proposed that the stochastic nature of sensory representations provide the means to implicitly encode sensory likelihoods ^81,82^. Others have shown that task-related firing rates of single neurons before the presentation of sensory information may be modulated by prior expectations ^83–85^, and firing rates after the presentation of sensory information may reflect Bayesian estimate of behaviorally-relevant variables ^30,86–88^. There have also been attempts to apply reliability-weighted linear updating schemes –commonly used in cue combination studies ^89–91^ – to explain how single-neuron firing rates might combine sensory evidence with prior expectations ^92,93^. However, the fact that single neurons encode various components of Bayesian models has not led to an overarching framework for understanding how networks of neurons perform Bayesian computations.

The central challenge in understanding Bayesian computations is the need for a framework that could bridge explanations at multiple scales. At one end are cellular-level explanations of how past experiences alter synaptic coupling between neurons, and on the other, are explanations of behavior based on the abstract notion of prior knowledge. This challenge was clearly stated by Nobel laureate Richard Axel ^94^, “we do not know the language by which […] patterns of neural activity are […] translated into appropriate behavioral or cognitive output.” In the case of Bayesian integration, we need a language that has the potential to explain how synaptic coupling between neurons could mediate prior-dependent biases in behavior.

Theoretical studies and recent artificial neural network models have established a framework that could potentially address this challenge. They indicate that structured connectivity creates low-dimensional activity patterns across the population with powerful computational capacities ^25,95^ for integration ^96^, categorization ^22^, gating ^97^, timing ^21,26,61,98^, learning ^99–101^, movement control ^80, 102–106^ and forming addressable memories ^107^. According to this framework, the key to a deeper understanding of how neural circuits perform computations is an analysis of the geometry and dynamics of activity across the population ^108^.

Using this approach, we found a novel computational principle for how neural circuits perform Bayesian integration. We found that prior statistics that were presumably embedded in the coupling between neurons, established low-dimensional curved manifolds across the population. This curvature, in turn, warped the underlying neural representations and afforded biases in accordance with Bayes-optimal behavior. This mechanism was evident across multiple behavioral conditions including different prior distributions and different effectors suggesting that it may be a general computational strategy for Bayesian integration.

Remarkably, this computational strategy also emerged spontaneously in an artificial neural network trained on the same sensorimotor task. Moreover, the network allowed us to probe the causal role of the underlying mechanisms *in-silico* using a set of targeted-dimensionality perturbation experiments that are currently not possible *in-vivo*. These experiments allowed us to reveal the role of bias in compensating uncertainty (i.e.. larger biases for noisier measurements), and validated the role of the curved manifold for integrating prior knowledge into behavioral responses. To investigate the neurobiological instantiation of Bayesian integration at the level of cells and synapses, future experiment should examine how functional and causal measures of coupling between neurons may change while such prior-dependent curved manifolds are formed across the population ^109^, as is done for simpler kinds of motor learning ^110^.

Although we focused on Bayesian integration in the domain of time, the key insights gleaned from our results are likely to apply more broadly to the general problem of Bayesian integration in perception, sensorimotor function and cognition. For example, numerous studies have found an important role for natural scene statistics in vision, and have further shown that the organization of tuning in neurons of the primary visual cortex follow those statistics ^8^. This observation is often explained in terms of efficient coding ^111^, which is a statement about the nature of the representation, and not about the underlying computations. We also found that single neurons developed flexible tuning for the range of intervals the animal was exposed to (Figure 2). In other words, single neurons in our experiment also abided by the principles of efficient coding. However, what distinguishes our approach is that it does not stop at the representational description. Instead, our results show how biased tuning across single neurons leads to warped representations in population dynamics whose geometry can explain the underlying Bayesian computations. We speculate that the same framework may provide valuable insights into the link between efficient coding and Bayesian perception ^112,113^, as well as numerous other sensorimotor and cognitive functions.

## Online Methods

All experimental procedures conformed to the guidelines of the National Institutes of Health and were approved by the Committee of Animal Care at the Massachusetts Institute of Technology. Experiments involved two awake, behaving monkeys (species: M. mulatta; ID: H and G; weight: 6.6 and 6.8 kg; age: 4 yrs old). Animals were head-restrained and seated comfortably in a dark and quiet room, and viewed stimuli on a 23-inch monitor (refresh rate: 60 Hz). Eye movements were registered by an infrared camera and sampled at 1kHz (Eyelink 1000, SR Research Ltd, Ontario, Canada). Hand movements were registered by a custom single-axis potentiometer-controlled joystick whose voltage output was sampled at 1kHz (PCIe6251, National Instruments, TX). The MWorks software package (http://mworks-project.org) was used to present stimuli and to register hand and eye position. Neurophysiology recordings were made by 1-3 24-channel laminar probes (V-probe, Plexon Inc., TX) through a bio-compatible cranial implant whose position was determined based on stereotaxic coordinates and structural MRI scan of the two animals. Signals were amplified, bandpass filtered, sampled at 30 kHz, and saved using the CerePlex data acquisition system (Blackrock Microsystems, UT). Spikes from single-units and multi-units were sorted offline using Kilosort software suites ^114^. Analysis of both behavioral and spiking data was performed using custom MATLAB code (Mathworks, MA).

### Two-prior time-interval reproduction task

Animals were trained on an interval-timing task that we refer to as the Ready-Set-Go (RSG) in which they had to measure a sample interval, *t_s_*, and produce a matching interval *t_p_* by initiating a saccade or by moving a joystick. Each trial began with the presentation of a circle (diameter: 0.5 deg) and a square (side: 0.5 degree) immediately below it. Animals had to fixate the circle and hold their gaze within 3.5 deg of it. The square instructed animals to move the joystick to the central location. To aid the hand fixation, we briefly presented a cursor whose instantaneous position was proportional to the joystick’s angle and removed it after successful hand fixation. Upon successful fixation and after a random delay (500 ms plus a random sample from an exponential distribution with mean of 250 ms), a white movement target was presented 10 deg to the left or right of the circle (diameter: 0.5 deg). After another random delay (250 ms plus a random sample from an exponential distribution with mean of 250 ms), the Ready and Set stimuli were flashed sequentially around the fixation cues (outer diameter: 2.2 deg; thickness: 0.1 deg; duration: 100 ms). The animal had to measure the sample interval, *t_s_*, demarcated by Ready and Set, and produce a matching interval, *t_p_*, after Set by making a saccade or by moving the joystick toward the movement target presented earlier (Go). Across trials, *t_s_* was sampled from one of two discrete uniform prior distributions each with 5 equidistant samples, a “Short” distribution between 480 and 800 ms, and a “Long” distribution between 800 and 1200 ms.

The full experiment consisted of 8 randomly interleaved conditions, 2 effectors (Hand and Eye), 2 movement targets (Left and Right), and two prior distributions (Long and Short). The 4 conditions associated with the 2 effector and 2 prior condition were interleaved randomly across blocks of trials. For 15 out of 17 sessions, the block size was set by a minimum (3 and 5 trials for H and G, respectively) plus a random sample from a geometric distribution with a mean of 3 trials, and was capped at a maximum (20 for H and 25 for G). The resulting mean ± SD block lengths were 4.0 ± 4.4 and 13.3 ±3.1 trials for H and G, respectively. In 2 sessions in H, switches occurred on a trial-by-trial basis. Because animal G had more trouble switching between conditions, block switches involved a change of prior or effector but not both. The position of the movement target was randomized on a trial-by-trial basis. Throughout every trial, the fixation cue provided information about the underlying prior and the desired effector. One of the two fixation cues was colored and the other one was white. The animal had to respond with the effector associated with the colored cue (circle for Eye and square for Hand) and the cue indicated the prior condition (red for Short and blue for Long).

To receive reward, animals had to move the desired effector in the correct direction, and the magnitude of the relative error defined as ∣*t_p_-t_s_*∣*/t_s_* had to be smaller than 0.15. When rewarded, reward decreased linearly with relative error, and the color of the response target changed to green. Otherwise, no reward was given and the target turned red. Trials were aborted when animals broke the eye or hand fixation prematurely before Set, used incorrect effector, moved opposite to the target direction, or did not respond within *3t_s_* after Set. To compensate for lower expected reward rate in the Long prior condition due to longer duration trials (i.e., longer *t_s_* values), we set the inter-trial intervals of the Short and Long conditions to 1220 ms and 500 ms, respectively.

## Behavior

We analyzed behavior in sessions with simultaneous neurophysiological recordings (H: 17 sessions, 26189 trials, G: 12 sessions, 30777 trials). First, we used a probabilistic mixture model to exclude outliers from further analysis. The model assumed that each *t_p_* was either a sample from a task-relevant Gaussian distribution or from a lapse distribution, which we modeled as uniform distribution extending from the time of Set to *3t_s_*. We fit the mean and standard deviation of the Gaussian for each unique combination of session, prior condition, *t_s_*, effector, and target directions. Using this model, we excluded any trial whose *t_p_* was more likely sampled from the lapse distribution (3.84% trials in H and 5.7% trials in G).

We measured the relationship between *t_p_* and *t_s_* separately for each combination of prior, effector, and target direction in individual sessions using linear regression (*t_p_=βt_s_*+*ε*). Since *t_p_* is more variable for larger *t_s_* due to scalar variability, we used a weighted regression - for each *t_s_*, error terms were normalized by the standard deviation of the distribution of *t_p_* for that *t_s_*. We tested whether regression slopes were larger than 0 and less than 1 (Figure 1, Figure S1, Table S1).

We also fit a Bayesian observer model to behavioral data (Figure 1, Figure S3). The Bayesian observer measures *t_s_* using a noisy measurement process that generates a variable measured interval, *t_m_*. The measurement noise has a Gaussian distribution with a mean of zero and a standard deviation that scales with *t_s_* with constant of proportionality *w_m_*. The observer combined the likelihood function, *p*(*t_m_*∣*t_s_*), with the prior, *p*(*t_s_*), and uses the mean of the posterior, *p*(*t_s_*∣*t_m_*), to compute an estimate, *t_e_*. The observer aims to produce *t_e_* through another noisy process generating a variable *t_p_*. We assumed that production noise scales with *t_e_* with constant of proportionality *w_p_*. For each prior, the model also included an offset term (*b*) to accommodate any overall bias in *t_p_*. Using maximum likelihood estimation, we fit the 4 free parameters of the model (*w_m_, w_p_, b_Short_*, and *b_Long_*) to data for each animal, effector, and target directions after pooling across sessions (Figure S3).

## Electrophysiology

We collected 456 single-units (H:196, G:260) and 902 multi-units (H:421, G:481) in 69 penetrations across 29 sessions (H:17, G:12). Most analyses were performed in a condition-specific fashion (2 priors, 5 *t_s_* per prior, 2 effectors, and 2 directions), and therefore, we excluded units for which we had less than 5 trials per condition. In addition, we excluded units whose average firing rate was less than 1 spike/s. The remaining units included in subsequent analyses were 536 and 636 in H and G, respectively.

We used a generalized linear model (GLM) to assess which neurons were sensitive to the prior and *t_s_*. We modeled spike counts in an 80-ms window immediately before Set, *r*_Set_, as a sample from a Poisson process whose rate was determined by a weighted sum of a binary indicator for prior (*l_prior_*: 1 for Long, 0 for Short) and 5 binary indicators for *t_s_* values associated with the Short prior for which we also knew the firing rate for the Long prior. The model was augmented by two additional binary indicators to account for independent influences of the effector (*l_effector_*: 1 for Hand, 0 for Eye), and direction (*l_direction_*: 1 for Left, 0 for Right).

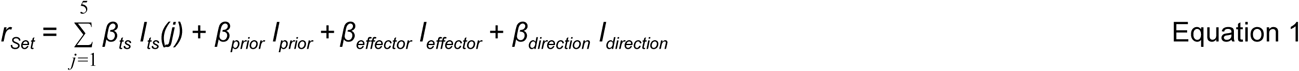

To get the most reliable estimate for the regression weights, we included spike counts based on all trials with attrition, and estimated *β* parameters of the model using MLE for all included neurons. To assess the significance of the effect of the prior condition, we used Bayesian information criteria (BIC) to compare the full model (Equation 1) to a reduced model that did not include a regressor for the prior (Equation 2):

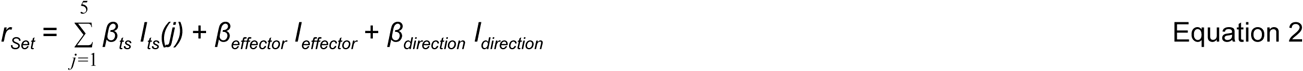

We also used a GLM to assess which neurons were sensitive to *t_s_*. Since values of *t_s_* were different between the priors, we used two distinct GLMs, one for data in the Short prior and one for the Long prior (Equation 3):

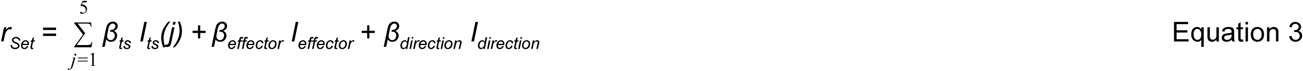

To identify the neurons that were sensitive to *t_s_*, we used BIC to compare the *t_s_*-dependent GLM (Equation 3) to a reduced GLM in which there was no sensitivity to *t_s_* (Equation 4):

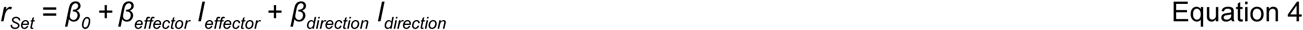

Neurons were considered *t_s_*-dependent if the BIC was lower in the full model either for the Short or for the Long prior condition (Figure 2).

## Population analysis

To examine the trajectory of population activity in state space, we applied principal component analysis (PCA) to condition-specific, trial-averaged firing rates (bin size: 20 ms, Gaussian smoothing kernel width: 40 ms). Since neurons modulated during estimation and production epochs were largely non-overlapping (Figure S6), we performed PCA separately on the two epochs. We first constructed firing rate matrices of all neurons and time points [time points x neurons]. This yielded 16 matrices (2 priors x 2 effectors x 2 directions x 2 epochs). We then concatenated the matrices across the two prior conditions along the time dimension and applied PCA to each of the resulting 8 data matrices to find principal components (PCs) for each unique combination of effector and direction, separately in the two epochs.

In the estimation epoch, firing rates for each *t_s_* were estimated with attrition (i.e., firing rate at time *t* was computed from spikes in all trials in which Set occurred after *t*). However, results were qualitatively unchanged if firing rates were estimated without attrition. In the production epoch, to accommodate different trial lengths (i.e., variable *t_p_*), we estimated firing rates only up to the shortest *t_p_* for each *t_s_*. Neural trajectories in the two epochs were analyzed within the subspace spanned by the top PCs that accounted for at least 75% of total variance (Figure S7). We will use ***X***(*t*) to refer to a neural state within the PC space at time *t*.

In the estimation epoch, we examined the rotational dynamics in neural trajectories during the support of each prior by projecting ***X***(*t*) onto an ‘encoding axis’, *u_ts_*, defined by a unit vector connecting the state associated with the shortest *t_s_* (*t_s_min_*) to that with the longest *t_s_* (*t_s_max_*) for that prior. We denote the projected states by ***X**u_ts_*. To reduce estimation error, we computed multiple difference vectors connecting ***X***(*t_s_min_*+*Δ*t) to ***X***(*t_s_max_-Δt*) for every *Δt*=20 ms, and used the average as our estimate of *u_ts_*. We used bootstrapping (resampling trials with replacement 1000 times) to compute 95% confidence interval for ***X**u_ts_*. We quantified the similarity between ***X**u_ts_* and the Bayesian estimates (*t_e_*) inferred from model fits to behavior using linear regression (***X**u_ts_* = *α* + *βt_e_*). Since we included spike counts across trials with attrition, there were nearly 5 times more data for the shortest *t_s_* compared to the longest *t_s_* within each prior. Accordingly, for each *t_s,_* error terms were weighted by the number of data points included for that t_s_. (5 for the shortest *t_s_*, 4 for the second shortest, and so forth). We then used the coefficient of determination (R^2^) to assess the degree to which *t_e_* was explained by the neurally inferred ***X**u_ts_*. Finally, we tested the specificity of our results with respect to the chosen *u_ts_* by performing the same analysis for 1000 randomly chosen encoding axes (*u’_ts_*), and comparing the corresponding R^2^ values.

In the production epoch, we defined **a** ‘decoding axis’, *v_tp_*, for each prior as the unit vector connecting the state associated with the shortest *t_s_* to that with the longest *t_s_* 200 ms after Set. We projected neural states 200 ms after Set onto *v_tp_* and compared the organization of projected states (***X**v_tp_*) to the Bayesian estimates (*t_e_*) using R^2^. We also performed the analysis for 1000 randomly chosen decoding axes (*v’_tp_*) to test the specificity of results with respect to the chosen *v_tp_*.

We also measured trial-by-trial correlation between ***X**u_ts_* and ***X**v_tp_* using a leave-one-out cross-validation procedure in one experimental session in animal H with a large number of simultaneously recorded neurons (N=48) and a large number of completed trials (1610 trials) (see Figure S10 for monkey G). For each condition (effector and direction), we computed PCs of trial-averaged firing rates across all neurons (including those recorded in other sessions) and all trials except the left-out trial.

We also analyzed neural activity at the level of single trials using cross-validation with the following procedure: (1) we designated one trial as test and the remaining trials as train dataset; (2) we binned and smoothed ***X*** for the test trial (20 ms for bin size and 40 ms for smoothing kernel size); (3) we projected the smoothed ***X*** onto *u_ts_* and *v_tp_* estimated from the train dataset to compute ***X**u_ts_* and ***X**v_tp_*. Repeating this procedure for different choices of test trial yielded distributions of ***X**u_ts_* and ***X**v_tp_* for individual trials. We then used the sensitivity index, d’ (i.e., difference between means relative to standard deviation) to quantify the distance of the distribution of ***X**u_ts_* for every *t_s_* to the distribution of ***X**u_ts_* for the mean *t_s_* for each prior condition. We also quantified the trial-by-trial correlation between ***X**u_ts_* and ***X**v_tp_*. To do so, we first standardized (i.e., z-scored) ***X**u_ts_* and ***X**v_tp_* values for each condition separately (2 priors, 5 *t_s_* values, 2 effector, and 2 target directions) and the combined the entire dataset to compute a reliable estimate of trial-by-trial correlations as well as 95% confidence interval derived from bootstrapping (Figure 4e). We repeated our measurement of correlation while using 1000 randomly chosen encoding and decoding axes (*u’_ts_* and *v’_tp_*) to further verify the validity of our choice of *u_ts_* and *v_tp_* (Figure 4f).

We finally examined two later links of the cascade model (Figure 3b) during the production epoch. A key component in the production epoch was the speed of the neural trajectory travelling the state space. For each dataset, we computed the speed as the average Euclidean distance (in the PC space accounting for at least 75% of the total variance) between neural states associated with successive bins (20 ms), divided by the duration separating Set+200ms and the time of Go. First, we related the trajectory speed to the projected state along the decoding axis (*v_tp_*) across the prior and *t_s_* to test if the state served as an initial condition to set up the speed of the ensuing trajectory (Figure 3G). We then assessed how the speed during the production epoch was associated with the behavioral output, *t_p_* (Figure 3H). We computed a correlation coefficient between the *t_p_* averaged across trials of each dataset and the trajectory speed and tested its statistical significance (p<0.05).

## Recurrent neural network

We constructed a randomly connected firing-rate recurrent neural network (RNN) model with N = 200 nonlinear units. The network dynamics were governed by the following equations:

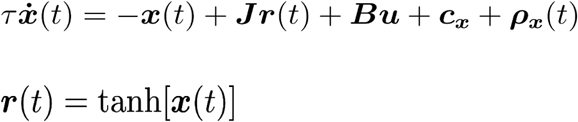

***x***(*t*) is a vector containing the activity of all units and ***r***(*t*) represents the firing rates of those units by transforming *x* through **a** `tanh` nonlinearity. Time *t* was sampled every millisecond for a duration of *T* = 3500 ms. The time constant of decay for each unit was set to *τ* = 10*ms*. The unit activations also contain an offset ***c**_x_* and white noise ***ρ**_x_*(*t*) at each time step with standard deviation in the range [0.01-0.015]. The matrix ***J*** represents recurrent connections in the network. The network received multi-dimensional input ***u*** through synaptic weights ***B*** = [***b**_c_,**b**_s_*]. The input ***u*** was comprised of a prior-dependent context cue *u_c_*(*t*) and an input *u_s_*(*t*) that provided Ready and Set pulses. In *u_s_*(*t*) Ready and Set were encoded as 20 ms pulses with a magnitude of 0.4 that were separated by time *t_m_*, which is the original interval *t_s_* transformed stochastically by weber noise *w_m_* (see next section for training details). The amplitude of the prior-dependent context input *u_c_*(*t*) was set to 0.3 for the short prior and 0.4 for the long prior contexts. Networks produced a one-dimensional output *z*(*t*) through summation of units with weights *w_o_* and a bias term *c_z_*.

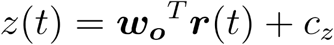

## Network Training

Prior to training, model parameters (*θ*), which comprised ***J, B, w_o_, c_x_*** and ***c_z_*** were initialized. Initial values of matrix ***J*** were drawn from a normal distribution with zero mean and variance 1/*N*, following previous work ^115^. Synaptic weights ***B*** = [***b**_c_, **b**_s_*] and the initial state vector ***x***(0) and unit biases ***c_x_*** were initialized to random values drawn from a uniform distribution with range [-1,1]. The output weights, ***w_o_*** and bias ***c_z_***, were initialized to zero. During training, model parameters were optimized by truncated Newton methods^116^ using backpropagation-through-time ^117^ by minimizing a squared loss function between the network output *z_i_*(*t*) and a target function *f_i_*(*t*), as defined by:

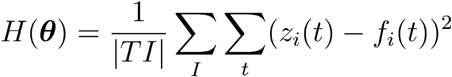

Here *i* indexes different trials in a training set (*I* = different prior contexts x intervals (*t_s_*) x repetitions (*r*)). The target function *f_i_*(*t*) was only defined in the production epoch (the output of the network was not constrained during the estimation epoch). The value of *f_i_*(*t*) was zero during the Set pulse. After Set, the target function was governed by two parameters that could be adjusted to make *f_i_*(*t*) nonlinear, scaling, non-scaling or approximately-linear:

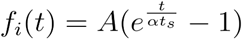

For the networks reported, *f_i_*(*t*) was an approximately-linear ramp function parametrized by *A* = 3 and *α* = 2.8. Solutions were robust with respect to the parametric variations of the target function (e.g., nonlinear and non-scaling target functions). In trained networks, the production time, *t_p_* was defined as the time between the Set pulse and when the output ramped to a fixed threshold (*z_i_* = 1).

During training, we employed three strategies to obtain robust solutions. ***ρ***(***t***) was drawn from a normal distribution with standard deviation 0.05. Furthermore, networks were trained and tested with a noisy measured interval (*t_m_*) that was generated from the interval *t_s_* plus interval-dependent noise with the constant of proportionality (*w_m_*=0.05), while fixing the objective itself to *t_s_*.

## Network causal experimentation

To evaluate the importance of the encoding axis on the behavior of the RNN at the time of Go, we performed a targeted perturbation experiment involving changes of the network state along the encoding axis shortly before Set, which we refer to as ‘re-encoding’. We systematically altered network states along the *u_ts_* 20 ms before the onset of Set and examined the consequences of this perturbation on behavior. To verify our approach, we first performed a control experiment in which the perturbation was expected to have no appreciable effect on behavior. Specifically, we re-encoded the network state for each trial of each *t_s_* to the expected state for that *t_s_* under no perturbation (n = 3000 trials per re-encoding). In this control experiment, perturbation had no effect on behavior (as expected) when we used a protocol in which (i) we allowed the network to stabilize for 10 ms after re-encoding (on the same order as the time constant of individual units in the RNN), and (ii) administered the Set pulse 10 ms after stabilization (Figure 5d). Having established a working protocol for the re-encoding experiment, we performed two causal experiments involving compression and translation of network states on *u_ts_*.

For the compression experiments, we evaluated the network’s behavior after applying various levels of compression (40% and 80%) to network states along *u_ts_* toward the mean state (i.e. the state associated with the mean of the prior). For the translation experiments, everything was the same except that the re-encoding involved a 20% shift in network states in the positive or negative directions (i.e., resulting in increasing or decreasing *t_s_*) (Figure 5e). One constraint in the translation experiment was that the network could not tolerate large negative shifts (i.e., intervals shorter than 400 ms for the short prior and 800 ms for the long prior). Such translations placed the network state in regions of the state space in which the latent dynamics were no longer governed by the rotating manifold.

## Acknowledgements

H.S. and N.M. are supported by the Center for Sensorimotor Neural Engineering. D.N. was supported by the Rubicon grant (446-14-008) by the Netherlands Scientific Organization and the Marie Sklodowska Curie Reintegration Grant (PredOpt 796577) by the European Union. M.J. is supported by NIH (NINDS-NS078127), the Sloan Foundation, the Klingenstein Foundation, the Simons Foundation, the McKnight Foundation, the Center for Sensorimotor Neural Engineering, and the McGovern Institute.

Author contributions
H.S. and M.J. conceived the *in-vivo* experiments. H.S. collected the physiology data. D.N. and M.J. conceived the *in-silico* experiments with recurrent neural networks. D.N. trained and simulated the networks. H.S., N.M. and D.N. analyzed the data. M.J. supervised the project. All authors were involved in writing the manuscript.

